# An Exclusively Skewed Distribution of Pediatric Immune Reconstitution Inflammatory Syndrome Towards the Female Sex is Associated with Advanced Acquired Immune Deficiency Syndrome

**DOI:** 10.1101/550129

**Authors:** Regina Célia de Souza Campos Fernandes, Thaís Louvain de Souza, Thiago da Silva Barcellos, Enrique Medina-Acosta

## Abstract

In human immunodeficiency virus and acquired immune deficiency syndrome (HIV/AIDS) patients with very low CD4 cell counts, there is a temporal relationship between administration of antiretroviral therapy (ART) and an increased inflammatory response state known as the immune reconstitution inflammatory syndrome (IRIS). The predominant clinical presentation of IRIS is an infectious disease that can be life-threatening. IRIS-related infectious events are distributed similarly between adult males and females, albeit a few studies have shown a skewing towards the male sex in pediatric IRIS. Here, we assessed sex-specific differences in the causes and extent of IRIS infectious events in HIV-infected pediatric patients on ART. We carried out a prospective clinical analysis (from 2000 to 2018) of IRIS-related infectious events after ART in a cohort of 82 Brazilian children and adolescents infected with HIV-1 through mother-to-child transmission as well as a comprehensive cross-referencing with public records on IRIS-related infectious causes in pediatric HIV/AIDS. Twelve events fulfilling the criteria of IRIS occurred exclusively in eleven females in our cohort. The median age at IRIS events was 3.6 years. The infectious causes included *Mycobacterium bovis*, varicella-zoster virus, molluscum contagiosum virus, human papillomavirus, cytomegalovirus, and *Mycobacterium tuberculosis*. In one female, there was regional bacillus Calmette-Guérin dissemination and cytomegalovirus esophagitis. There was complete health recovery after ten IRIS events without the use of corticosteroids or ART interruption. One case of IRIS-associated miliary tuberculosis was fatal. The biological female sex was a significant risk factor for IRIS events (odds ratio: 23.67; 95% confidence interval 95%: 1.341 – 417.7; *P* = 0.0016). We observed an effect of the advanced HIV/AIDS variable in IRIS females as compared with non-IRIS females (mean CD4+ T cell percentage 13.36% versus 18.63%; *P* = 0.0489), underpinning the exclusively skewed distribution towards the female sex of this cohort. Moreover, the IRIS females in our cohort had higher mean CD4^+^ T cell percentages before (13.36%) and after IRIS (26.56%) than those of the IRIS females (before IRIS, 4.978%; after IRIS, 13.81%) in previous studies conducted worldwide. We concluded that the exclusively skewed distribution of pediatric IRIS towards females is associated with more advanced AIDS.

## 1. Introduction

Rigorous adherence to antiretroviral therapy (ART) leads to recovery from immunodeficiency and results in a rapid decrease in morbidity and mortality rates among human immunodeficiency virus (HIV)-1 infected patients. In ART patients with very low CD4 cell percentages (CD4%), there is a temporal relationship between therapy and an increased inflammatory response state known as the immune reconstitution inflammatory syndrome (IRIS), occurring a few weeks to months after therapy administration. The onset of IRIS involves clinical manifestations that can be life-threatening and coincides with an elevation in CD4% and a drop in HIV-1 loads (French et al., 2004; Stoll and Schmidt, 2004; French, 2009). In most cases, IRIS manifests as opportunistic infections. IRIS-related infectious events can be classified as unmasking, in which there is a subclinical and therefore unrecognized infection that is unveiled after ART, or paradoxical, in which there is an exacerbation of an infectious disease previously observed in the patients (French, 2009).

The most common infectious agents associated with IRIS manifestations are tuberculous (TB) or nontuberculous mycobacteria, cryptococci, herpesvirus, cytomegalovirus (CMV), hepatitis B and C viruses, John Cunningham virus, and Pneumocystis spp. (French et al., 2004). In severely immunocompromised HIV-infected adults, the onset of IRIS-related TB ranges from 10 to 14 days after ART (Vignesh et al., 2013). In pediatric HIV/AIDS, IRIS-related TB can occur up to six months after ART (Narendran et al., 2006; Puthanakit et al., 2006b; Hatherill and Flisher, 2009; Innes et al., 2009; Wang et al., 2009; de Carvalho et al., 2010; Rabie et al., 2010; Kalk et al., 2013). Grave’s autoimmune disease as a manifestation of late-onset IRIS has been reported in adults (Rasul et al., 2011), but in only one pediatric patient (Perez et al., 2009). When corticosteroid treatment of IRIS-associated infections is ineffective and life is threatened (Safdar et al., 2002), ART interruption must be considered (Shah, 2011).

IRIS-related infectious events remain a challenge in the managing of HIV-infected pediatric patients. Notwithstanding the scarcity of studies, the incidence of IRIS in children on ART ranges from 4.7 to 38% (Orikiiriza et al., 2010; Walters et al., 2014) and the associated risk factors vary from one study to another. For example, in one study on 162 pediatric patients from Uganda, male sex, pretreatment low CD4%, CD8 cell count, and coughing were determined IRIS risk factors (Orikiiriza et al., 2010). In a second study on 494 pediatric patients from South Africa (Walters et al., 2014), increased risk for IRIS was associated with age < 12 months. In another study, viral loads in the IRIS group were found significantly higher than in the control group (Wang et al., 2009), whereas lower CD4% in subjects presenting with IRIS-related infectious events was a risk factor (Puthanakit et al., 2006a).

Although a statistically significant association between a biased male/female sex ratio and IRIS has not described (Shah, 2011), a trend towards the male sex has been reported (Orikiiriza et al., 2010). Herein, we assessed sex-specific differences in the causes and extent of IRIS-related infectious events in HIV-infected Brazilian pediatric patients on ART.

## 2. Materials and Methods

### 2.1 Study design and experimental setting

We prospectively observed children and adolescents with confirmed HIV infection and on ART (n = 82) from August 2000 to August 2018 at the Specialized Assistance Service of the Municipal Program for the Surveillance of Sexually Transmitted Diseases and AIDS of the city of Campos dos Goytacazes (population 463,731; 2010 census), Rio de Janeiro, Brazil. The cohort includes two cases of IRIS-associated BCGitis, whose clinical presentations were previously reported (Fernandes et al., 2009; Fernandes and Medina-Acosta, 2010).

### 2.2 Ethical considerations

The study received approval (FR-405294) from the Regional Committee of Ethics in Research in Humans from the Faculty of Medicine of Campos. All legally authorized next-of-kin gave written informed consent on behalf of participants in compliance with the Declaration of Helsinki. Clinical examination was performed by only one infectious disease specialist pediatrician (RC). The follow-up was performed monthly or whenever the clinical condition demanded.

### 2.3 Antiretroviral therapy

ART was provided universally for infants with confirmed HIV infection during the first year of life and for children or adolescents with moderate or severe clinical manifestations or immunodepression (CD4% < 25%), following the recommendations of the Brazilian Ministry of Health (Brazil, 2009). It is of note that during the 18 years of the study, different ART regimens were implemented according to the national treatment guidelines.

### 2.4 Definition criteria of IRIS

We used the major and minor criteria for IRIS listed by French and colleagues (French et al., 2004); cases required compliance with the two major criteria or one major criterion plus two minor criteria for inclusion. The major criteria are (i) atypical presentation of opportunistic infections or tumors in patients responding to ART and (ii) decrease in plasma HIV RNA concentration by > 1 log copies/mL. The minor criteria are (i) increase in blood CD4% after ART, (ii) increase in an immune response specific to the relevant pathogen, and (*iii*) spontaneous resolution of the infectious episode without specific antimicrobial therapy or tumor chemotherapy with the continuation of ART (French et al., 2004).

### 2.5 Estimates of X-chromosome inactivation

Genomic DNA samples were extracted from peripheral blood of females with IRIS-related infectious events. The extent of X-chromosome inactivation (XCI) was then estimated by interrogating the 5^me^CpG epigenetic marks neighboring short tandem repeats localized in the promoter regions of either the X-linked retinitis pigmentosa *RP2* gene (Xp region) or the androgen receptor *AR* gene (Xq region), as we had previously reported (Machado et al., 2014).

### 2.6 Searching relevant cases via PubMed

We carried out a literature review following the Preferred Reporting Items for Systematic Reviews and Meta-Analyses (PRISMA) statement guidelines (Moher et al., 2009) to extract data regarding IRIS-related events from clinical case reports and case series published in the English language. We searched by relevant biomedical tags in the PubMed database of the National Library of Medicine (https://www.ncbi.nlm.nih.gov/pubmed/) from January 1^st^, 1979 through August 30^th^, 2018 using the EndNote X9 (Clarivate Analytics, Philadelphia, PA) reference managing software. The following terms were used in pairs, immune reconstitution inflammatory syndrome, IRIS, HIV, child, children, immunodeficiency, and infant. To be manually reviewed by two annotators and listed as relevant semantic context, a clinical case was required to be a child or adolescent (age < 18 years old), have at least a positive HIV-1 serologic test, and an opportunistic infection with atypical presentation after ART introduction.

### 2.7 Data analysis

We used the EpiInfo^™^ public suite from the Centers for Disease Control and Prevention, USA (Christiansen and Lauritsen, 2010) to analyze data variables and to statistically evaluate possible associations between risk factors and the observed IRIS outcome.

## 3. Results

Throughout 18 years, we prospectively studied 82 HIV-infected children and adolescents on ART (36 males and 46 females). Two patients, male and female, were lost to specialist follow-up because they moved to another city or country. In the remaining, no males presented with IRIS, while 11 (13.8%) females developed 12 IRIS-related infectious events (risk ratio [RR]: 23.67; 95% confidence interval [CI]: 1.341–417.7; *p* = 0.0016). The median time to first symptom presentation following ART was 60 days (mean 108.6 ± 42.41 days). Clinical manifestations, laboratory findings, therapeutic approaches, and evolution of patients with IRIS-related infectious events are summarized in Table 1. For subjects that did not present with IRIS-related events, laboratory findings are listed in Supplementary Table S1.

**Table 1.** Characteristics of clinical manifestations and staging, laboratory findings, therapeutic approach, and evolution of patients with IRIS.

| Patient | Characteristics of HIV-infection |  |  |  |  |  | Characteristics of IRIS events |  |  |  |  |
| --- | --- | --- | --- | --- | --- | --- | --- | --- | --- | --- | --- |
|  | ART regimen <sup>a</sup> | Stage | CD4 (%) |  | Viral load (copies/mL) |  | Age at IRIS | Year of event | ART timing <sup>a</sup> | Clinical manifestation | Evolution |
|  |  |  | Basal | IRIS | Basal | IRIS |  |  |  |  |  |
| 1 | DDI, 3TC, EFV, LPV/Rt | A3 | 2.4 | 25.7 | 19,000 | 1,400 | 12 years | 2003 | 6 months | Genital HPV | complete recovery |
| 2 | AZT, DDI, NVP | A3 | 9.6 | 30.4 | 110,000 | 990 | 4 years | 2004 | 3 months | Molluscum contagiosum | complete recovery |
| 3 | AZT, 3TC, LPV/Rt | C3 | 13.3 | 29.7 | 5,130,000 | 1,410 | 8 months | 2006 | 41 days | Regional BCG dissemination | complete recovery |
| 4 | AZT, 3TC, LPV/Rt | C1 | 27.2 | 30 | 1,200,000 | 11,400 | 7 months | 2006 | 19 days | Regional BCG dissemination | complete recovery |
| 4 | AZT, 3TC, LPV/Rt | C1 | 27.2 | 30 | 1,200,000 | 11,400 | 7 months | 2006 | 2 months | Esophagus disease by CMV | distal esophageal stenosis |
| 5 | AZT, 3TC, EFV | A3 | 9 | 20 | 137,373 | 689 | 7 years | 2008 | 20 days | Herpes Zoster | complete recovery |
| 6 | AZT, 3TC, LPV/Rt | C3 | 9.9 | 21 | 62,300 | undetected | 3 years | 2008 | 18 months | Perianal HPV | complete recovery |
| 7 | AZT, 3TC, EFV | C3 | 13.47 | 24.45 | 334,545 | 803 | 21 months | 2010 | 2 months | Molluscum contagiosum | complete recovery |
| 8 | TDF, 3TC, EFV, DRV, Rt | C3 | 19.79 | 25.49 | 20,272 | undetected | 18 years | 2011 | 4 months | Herpes Zoster | complete recovery |
| 9 | AZT, 3TC, EFV | A3 | 8.2 | 8.97 | >500,000 | 937 | 9 years | 2012 | 5 months | Herpes Zoster | complete recovery |
| 10 | TDF, 3TC, LPV/Rt | C3 | 14.29 | – | 1,012,485 | – | 16 years | 2014 | 14 days | Miliary tuberculosis | death |
| 11 | AZT, 3TC, LPV/Rt | C3 | 6.01 | 46.40 | 48,519 | 4,473 | 10 years | 2016 | 9 days | Tuberculous pneumonia | complete recovery |

<sup>a</sup> DDI (Didanosine); 3TC (lamivudine); EFV (Efavirenz); LPV (Lopinavir); Rt (Ritonavir); NVP (Nevirapine); DRV (Darunavir); TDF (Tenofovir)
<sup>b</sup> Time to onset of IRIS event after ART

### 3.1 Clinical summary

There were two *Mycobacterium bovis* bacillus Calmette-Guérin (BCG)-related IRIS infectious events 41 and 18 days after ART introduction; they were treated with isoniazid (10 mg/kg/day) and ART was maintained. Surgical manipulation of lesions was contraindicated. Varicella-zoster virus occurred in three cases of dermatomal disease with excellent response to acyclovir therapy. Human papillomavirus infection was implicated in two IRIS cases. Two more IRIS cases were caused by molluscipox infection in the thoracic region with good evolution. One infant female presented with ulcerations at the posterior palate two weeks before ART that was managed with acyclovir; she developed vomiting and feeding intolerance and was diagnosed with an esophageal stricture. CMV exposure was confirmed by serology at 12 months (immunoglobulin [Ig] G: 1070.7 UA/mL; IgM: 0.86 UA/mL). Exteriorization of the proximal esophagus, gastrostomy, and dilatation were performed and in 2017, her digestive tract was successfully reconstructed. An adolescent female, without adequate adherence to ART and with pulmonary TB that was treated earlier, presented with weight loss, cervical adenopathies, respiratory distress, and miliary radiological pattern after 14 days of supervised ART, which progressed to death in three days. Lastly, a 10-year-old female was diagnosed with bacterial pneumonia nine days after commencing ART. At home, she was treated with penicillin and developed fever, respiratory distress, weight loss, and bilateral lung infiltrates after 60 days, requiring hospitalization. She had a negative tuberculin test result, albeit household TB contact was reported. She was then treated with rifampicin, isoniazid, and pyrazinamide with complete recovery.

In total, we observed 11 episodes of unmasked IRIS and one of paradoxical IRIS. When comparing viral loads (Figure 1A, B) and CD4% (Figure 1C, D), females with IRIS had significantly (P = 0.0489) lower CD4 values than those of non-IRIS females and viral loads were similar to those of non-IRIS females and males (Figure 1).

**Figure 1.**
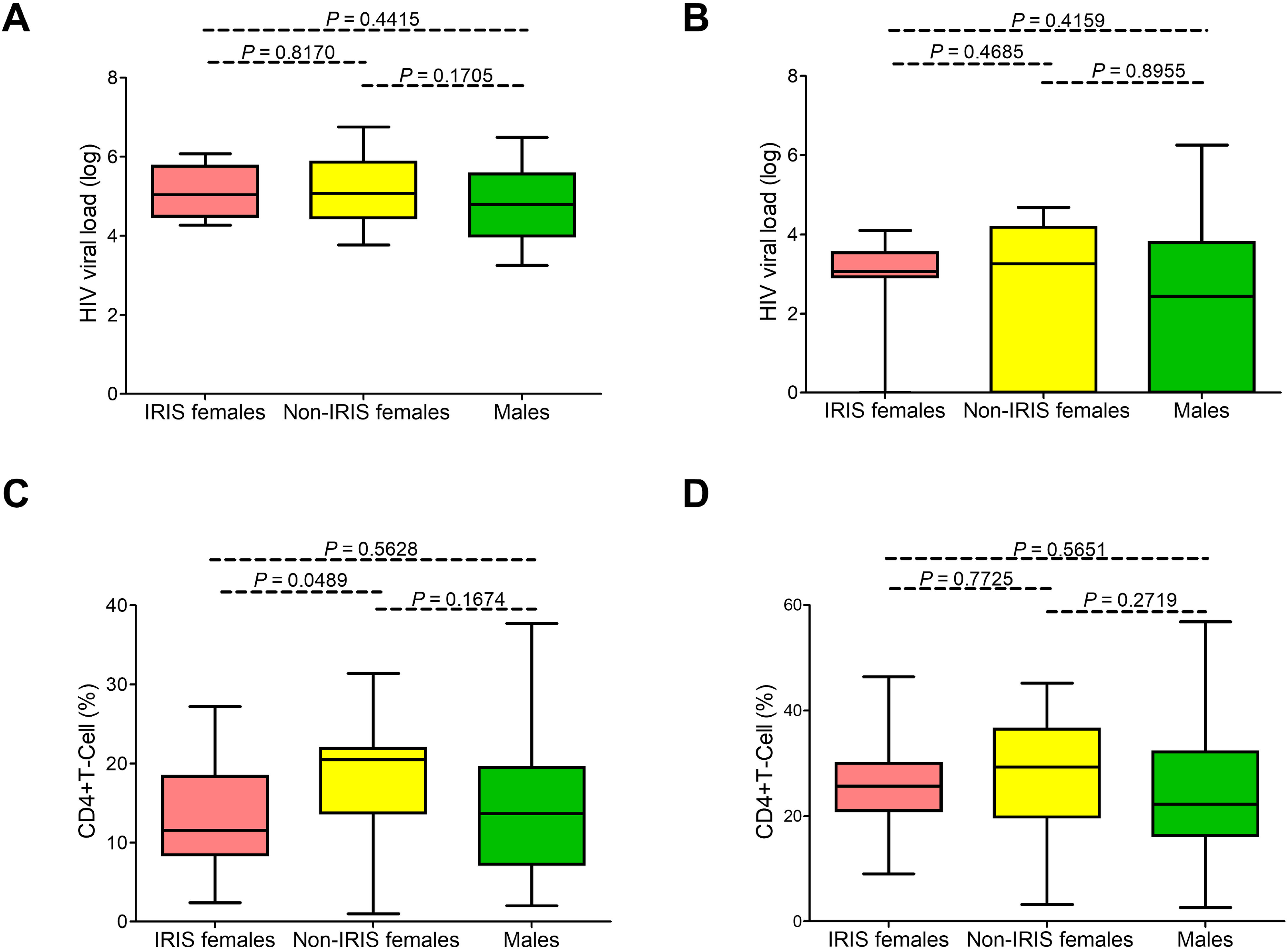
Virologic and immunology profiles in the Brazilian pediatric HIV cohort. No statistical difference was observed between viral loads before (**A**) and after ART (**B**) in females presenting with IRIS-related infectious events (represented in pink), females with no IRIS events (yellow), and males (green). Females presenting with IRIS infectious events exhibited lower CD4 cell percentages before (**C**) but not after ART (**D**).

The exclusively skewed occurrence of IRIS-related infectious events towards pediatric females is intriguing. Therefore, we investigated a possible association between extremely skewed female sex distribution of IRIS manifestations and affected females exhibiting highly skewed XCI. We only had genomic DNA samples available from six female patients for the XCI assay. Notably, only one affected female exhibited a preferential (> 90%) XCI (Supplementary Figure S1).

### 3.2 IRIS-related infectious events in pediatric HIV/AIDS reported in the literature

The literature search yielded 169 publications, but only 40 studies passed our requirements for listing relevant cases (Supplementary Table S2). In total, we listed 127 HIV-1 infected children and adolescents with IRIS-related infectious events. Most cases (80.1%) were from Peru, South Africa, and Thailand. IRIS-related events were distributed between males (37.8%) and females (48%); gender was not indicated for 18 reported cases. The age at IRIS-related infectious events exhibited a bimodal distribution with 28.3% (36/127) of cases occurring before age 1-year and 32.3% (41/127) between the ages of 8- to 11-year. The period from commencing ART to IRIS manifestation ranged from 0 to 120 days, with 54.3% (69/127) of cases occurring six weeks after ART. Fatal outcomes after IRIS-related disease occurred in 12.6% (16/127) of cases.

Out of all IRIS-related events, 14.2% (18/127) were pulmonary tuberculosis (Supplementary Table S2), 7.0% (9/127) nontuberculous mycobacterial disease, and 22.8% (29/127) BCG vaccine adverse events. Neurological disorders like progressive multifocal leukoencephalitis and cryptococcal meningitis occurred in 2.4% (3/127) and 3.1% (4/127) of the cases, respectively.

Viral load >1000 copies (> 3 log) before IRIS events was observed in 84.9% (79/93) of cases. After the onset of IRIS-related infection events, viral loads < 1000 (< 3 log) were reported in 70.6% (60/85) of cases, with 44.9% presenting with >1000 (> 3 log); in 11.8% (10/85) of cases, the viral load was undetectable. When restricting the analysis to female subsets from the compilation and our cohort, the viral loads were found decreased in both subgroups (Figure 2A, B).

**Figure 2.**
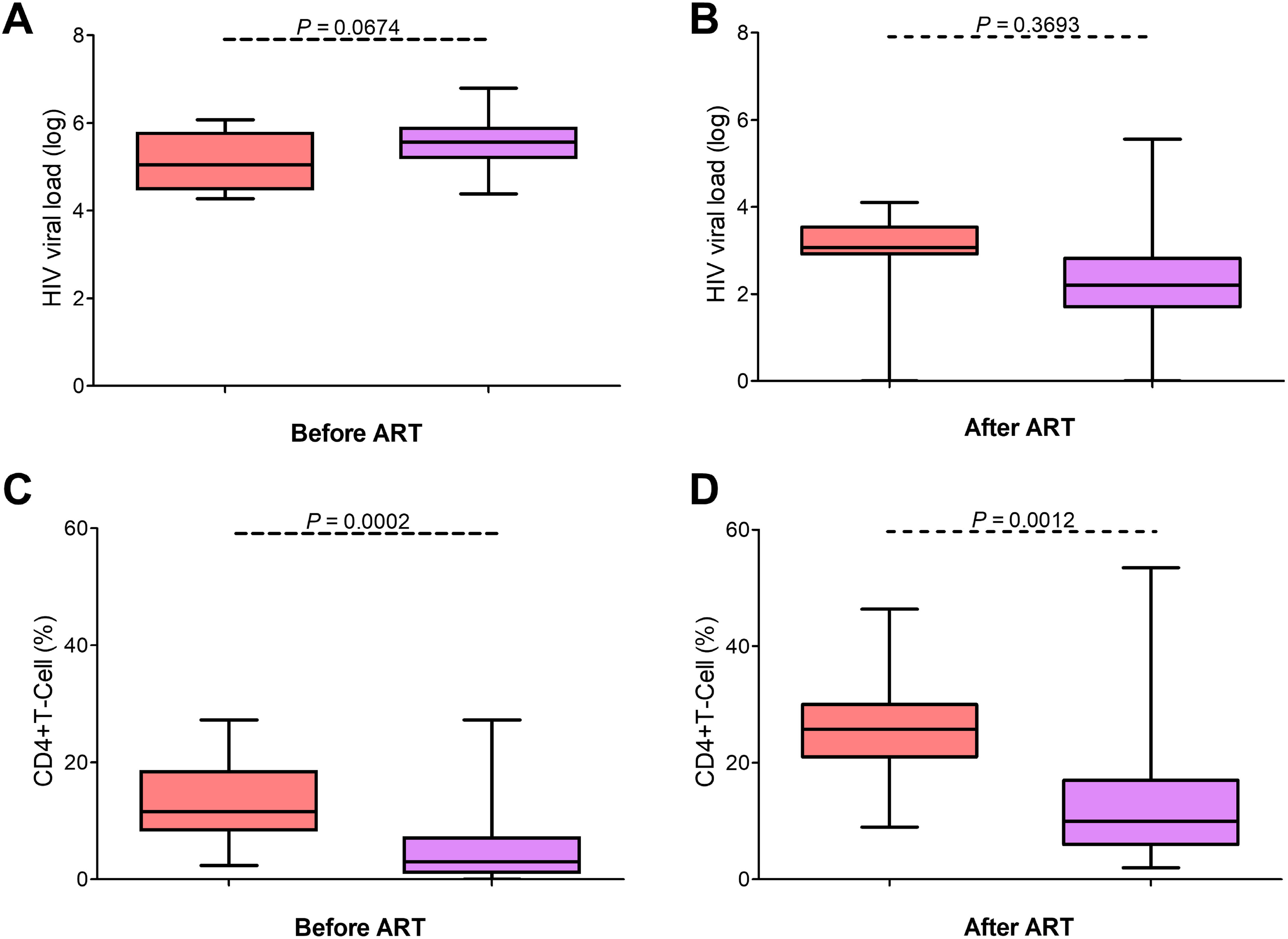
Cross-referencing virologic and immunology profiles in IRIS patients with published reports. Compared with pediatric HIV female patients presenting with IRIS-related infectious events listed in the literature compilation (represented in lilac), females with IRIS in our cohort (pink) exhibited lower viral loads (**A**) and higher CD4 cell percentages after ART (**D**). No statistical difference was observed between the viral loads in our cohort (**B**; pink) and the cases reported in the literature compilation (**C**; blue). Females presenting with IRIS infectious events in the literature compilation presented lower CD4 cell percentages (**E**) than the females with IRIS in our cohort (**F**).

Before the onset of IRIS, 85.7% (96/112) of the reported cases had CD4 cell counts < 200 cells/mL (< 15%). After IRIS, CD4 cell counts increased to 200–499 (> 15%) in 40.4% (40/99) of the cases. When the female subsets were compared, we noted that the compilation subset had significantly lower CD4 counts (P = 0.0002) than that of our cohort and that there was an increase in CD4% in both groups (Figure 2C, D). The IRIS females of our cohort had higher mean CD4+ T cell percentages before (13.36%) and after IRIS (26.56%) than those of the IRIS females (before IRIS, 4.9%; after IRIS, 13.8%) in previous studies conducted worldwide. Interestingly, the IRIS females in the literature have more advanced HIV/AIDS than the IRIS females of our cohort.

## 4. Discussion

We identified an abnormal distribution, fully skewed towards the female sex, of IRIS-related infectious events in HIV-infected Brazilian children and adolescents on ART. We observed 12 episodes in 11 females and none in males during a study period of 18 years. The bulk of clinical data allowed us to suggest that the observed skewed distribution towards the female sex is due to more advanced HIV/AIDS in these females than in non-IRIS females. Cross-referencing with pediatric HIV/AIDS data from the literature revealed a fair distribution of reported IRIS-related infectious events between females and males (Puthanakit et al., 2006a; Wang et al., 2009; Shah, 2011; Walters et al., 2014), except for a study in Uganda that reported a bias towards the male sex (P = 0.010) (Orikiiriza et al., 2010). In adults with HIV/AIDS, IRIS-related infectious events occurred at the same rate in both females and males (Breton et al., 2004; Murdoch et al., 2008; Kumar et al., 2012; He et al., 2013), albeit one study reported a male bias (P = 0.018; (Shelburne et al., 2005)). The IRIS females in our cohort had higher mean CD4+ T cell percentages before and after IRIS than those of the IRIS females in previous studies conducted worldwide.

Sex-biased susceptibility to bacterial infections has been linked to the differential effects of sex steroid hormones (estrogen and testosterone) on innate immunity (Garcia-Gomez et al., 2013); for example, males are more susceptible to TB than females (Stival et al., 2014). A caveat against sex hormones being involved in female-biased presentation of IRIS in our cohort is the fact that none of the males presented with IRIS. Unfortunately, we did not measure sex hormones in the 11 affected females at the time of IRIS events and it is thus unclear whether abnormal sex hormone levels are implicated in female-biased IRIS manifestations. We did investigate, however, whether this skewed IRIS event distribution towards females was associated with highly skewed XCI. A significant number of immune-associated genes map to the X-chromosome (Bianchi et al., 2012) and preferential XCI in females can result in phenotypic susceptibility and disease. In female eutherian mammals with normal XCI, the epigenetic transcriptional silencing of an X-chromosome in each somatic cell occurs at random during the early stages of embryonic development, assuring monoallelic expression in each cell and compensating for dosage-sensitive X-linked genes between females (XX) and males (XY) (Machado et al., 2014). Although we only had genomic DNA samples from six female patients, the complete skewing towards the female sex could not be explained by discrete differences in the rates of XCI tested in blood.

Our study exemplified the broad spectrum of etiological agents associated with IRIS-related infectious events in childhood and adolescence. IRIS-related BCG regional adenitis occurred in 16.6% (2/12) of cases, highlighting the breadth of association between HIV infection and BCG vaccination at birth (Puthanakit et al., 2005; Fernandes et al., 2009; Rabie et al., 2011). Esophageal strictures infrequently complicate the presentation of CMV disease in HIV-infected adults (Wilcox, 1999) and there appears to be a functional association between IRIS and esophageal stricture observed in CMV infection cases (Wilcox, 1999). To our knowledge, we report the first pediatric case of esophageal stricture secondary to CMV infection related to IRIS; moreover, CD4% during the IRIS event was > 25, corroborating the view that CD4% is not a reliable marker for disease progression and severity in infants.

IRIS-related infectious events should be considered important contributors to higher mortality rates in resource-limited settings with a late introduction of ART (Davies and Meintjes, 2009). Very early ART administration is an essential preventive factor against an IRIS-related fatal outcome (Rabie et al., 2011). Furthermore, IRIS-related infectious events are more life-threatening at an early age (Smith et al., 2009). In our cohort, six IRIS-related episodes occurred at less than one year of age. None of the 11 females were treated with corticosteroids and all remained on ART. Recovery was completed in 10 females, but there was a fatal case for a 16-year-old patient (mortality of 8.3%).

The prevalence of reported IRIS-related infectious events varied greatly by country or geographical region; 4.7% of the cases were reported in South Africa (Walters et al., 2014), 5.9% in the United Kingdom (Gkentzi et al., 2014), 11.5–16.4% in the USA (Tangsinmankong et al., 2004; Nesheim et al., 2013), 18.9% in India (Shah, 2011), 19% in Thailand (Puthanakit et al., 2006a), 20% in Peru (Wang et al., 2009), 38.3% in Uganda (Orikiiriza et al., 2010), 23% in Latin America (Krauss et al., 2015), and 22% in Malawi and Botswana (Cox et al., 2013). The lower rates may in part be explained by socio-economic status (i.e., better nutritional status of patients), moderate manifestations of HIV infection, and varying compliances with the definitions criteria. It is unclear whether differential (epi)genetic components can partly account for the disparity in distribution. Sex-stratified genome-wide association studies of IRIS using multiethnic genotyping arrays are needed to appraise the differences in disease susceptibility and to identify candidate autosomal and X-linked loci in diverse and admixed populations.

In conclusion, our prospective study found that IRIS-related infectious events occurred exclusively in females in a cohort of 80 HIV-infected Brazilian children and adolescents on ART. This complete skewing towards the female sex is uncommon and was linked to more advanced HIV/AIDS.

## Supporting information

Supplementary material full

## 5. Conflict of Interest

The authors declare that the research was conducted in the absence of any commercial or financial relationships that could be construed as a potential conflict of interest.

## 6. Author Contributions

RF, TS, EM: designed the study, analyzed data, wrote and edited the typescript. RF: performed the clinical follow-up. TS: performed XCI assays. TS, TB: carried out the literature review. All the authors gave final approval.

## 7. Funding

The study was supported by Conselho Nacional de Desenvolvimento Científico e Tecnológico – CNPq, Brazil (http://cnpq.br/) [grant number 301034/2012-5 and 308780/2015-9 to EM].

## 8. Acknowledgments

We thank all participants and their guardians for participating in this study.

## Supplementary Materials

**Table S1.** Clinical staging, classification, and monitoring characteristics for HIV/AIDS pediatric patients who did not present with IRIS-related infectious events in our cohort.

**Table S2.** PubMed-based compilation of IRIS-related infectious events in children and adolescents infected with HIV (January 1^st^, 1979 through August 30^th^, 2018).

**Figure S1.** Rates of X-chromosome inactivation (XCI) in six females presenting with IRIS-related infectious events. The percentage of XCI was determined by genotyping genomic DNA with the two highly polymorphic short tandem repeat loci located in the *AR* and *RP2* genes (Machado et al., 2014) to identify heterozygotes. Both *AR* and *RP2* genes escape XCI and thus in females with random XCI, the rate of X-chromosome active (Xa) over X-chromosome inactive (Xi) is approximately 50%. The figure depicts the Xa/Xi rate (in percentage) observed for the *AR* and *RP2* marker systems. Only one female sample exhibited a highly skewed rate of XCI (> 90%).

