## Supplementary material full for "An Exclusively Skewed Distribution of Pediatric Immune Reconstitution Inflammatory Syndrome Towards the Female Sex is Associated with Advanced Acquired Immune Deficiency Syndrome"

**Table S1**. Clinical staging, classification, and monitoring characteristics for HIV/AIDS pediatric patients who did not present with IRIS-related infectious events in our cohort.

| | | | Before ART | | After ART | |
|---|---|---|---|---|---|---|
| **Patient** | **Sex** | **Clinical staging** | **Viral load (log)** | **CD4 (%)** | **Viral load (log)** | **CD4 (%)** |
| 1 | Female | N | 5886 | – | 3494 | 27.26 |
| 2 | Female | C | 6739 | – | 4688 | – |
| 3 | Female | C3 | 4.1 | 1 | Undetectable | 3.2 |
| 4 | Female | C2 | – | 17.8 | Undetectable | 24 |
| 5 | Female | B3 | 4.95 | 13.7 | – | 16 |
| 6 | Female | C2 | – | – | 3681 | 34.04 |
| 7 | Female | B2 | – | 21 | 3207 | 25 |
| 8 | Female | A2 | – | 22 | Undetectable | 32.36 |
| 9 | Female | B2 | 5177 | 20.9 | – | 8 |
| 10 | Female | B2 | 3919 | 19.3 | Undetectable | 29.31 |
| 11 | Female | A2 | 4.45 | 22.6 | 3307 | 37.8 |
| 12 | Female | A | 5342 | – | Undetectable | 31.29 |
| 13 | Female | N2 | 5.25 | – | Undetectable | 37 |
| 14 | Female | N2 | 4442 | 20.98 | Undetectable | 41.02 |
| 15 | Female | N | 6581 | – | 2.91 | 29.22 |
| 16 | Female | A2 | 4261 | 21.94 | Undetectable | 40.7 |
| 17 | Female | N | – | – | 3.81 | 36.75 |
| 18 | Female | C3 | 5.07 | 11.2 | Undetectable | 19.8 |
| 19 | Female | B3 | 4778 | 13.7 | 4176 | 15.8 |
| 20 | Female | C3 | 5.9 | 12.6 | 3591 | 15.1 |
| 21 | Female | A2 | – | 21.92 | Undetectable | 28.26 |
| 22 | Female | B2 | 5863 | 20.8 | 2146 | 31.7 |
| 23 | Female | C2 | – | 28.4 | Undetectable | 45.2 |
| 24 | Female | C3 | 5.62 | – | 4.28 | 36.5 |
| 25 | Female | B3 | 4.74 | 13.8 | 2.3 | 16.2 |
| 26 | Female | B | 6.11 | – | 4477 | – |
| 27 | Female | N1 | 4244 | 31.4 | 3661 | 30.41 |

| | | | | | | |
|---|---|---|---|---|---|---|
| 28 | Female | C1 | 4.96 | – | 4.6 | – |
| 29 | Female | A3 | 5.49 | – | 4.58 | – |
| 30 | Female | N2 | 3771 | 20.2 | Undetectable | 29.8 |
| 31 | Female | C | 5.08 | – | 4.2 | – |
| 32 | Female | N2 | 4832 | 17.28 | Undetectable | 22.92 |
| 33 | Female | B | – | – | 3.9 | – |
| 34 | Female | C | 6754 | – | 4335 | – |
| 35 | Female * | N2 | – | 21.94 | – | 16 |
| 36 | Male | N | 5087 | – | 2.38 | 21.81 |
| 37 | Male | N | 5.65 | – | 3908 | 24.79 |
| 38 | Male | A | 5482 | – | 3647 | 31.47 |
| 39 | Male | B1 | 5682 | 25.59 | 5624 | 27.39 |
| 40 | Male | B1 | 4.8 | 37.7 | 2.17 | 56.8 |
| 41 | Male | A3 | 4.17 | 7.2 | Undetectable | 25.1 |
| 42 | Male | N2 | – | 23 | – | 32 |
| 43 | Male | A3 | – | 13.7 | Undetectable | 20 |
| 44 | Male | C1 | – | – | Undetectable | 7 |
| 45 | Male | N1 | 5361 | 37.34 | Undetectable | 38 |
| 46 | Male | N2 | 4088 | 19.72 | Undetectable | 25.3 |
| 47 | Male | B2 | 4591 | 21.3 | 3805 | – |
| 48 | Male * | NI | – | – | – | – |
| 49 | Male | B3 | 4.55 | 12.27 | 4265 | 18.5 |
| 50 | Male | A3 | 4.7 | 2 | 1.3 | 22 |
| 51 | Male | C3 | 4794 | 2.97 | 3718 | 2.64 |
| 52 | Male | C3 | – | 6.9 | Undetectable | 35.6 |
| 53 | Male | C3 | 6.25 | – | 5 | – |
| 54 | Male | N2 | 4 | 18 | 3892 | 32.2 |
| 55 | Male | B1 | 3.91 | – | 2.9 | – |
| 56 | Male | N2 | 3466 | 19.4 | Undetectable | 21.1 |
| 57 | Male | A2 | – | 11.4 | 2633 | 11.2 |
| 58 | Male | C3 | – | – | 3.75 | – |
| 59 | Male | A2 | 4.8 | – | Undetectable | – |
| 60 | Male | N3 | 3.53 | 12.2 | 2519 | 19.4 |
| 61 | Male | A3 | 3255 | 8.1 | – | 15.5 |
| 62 | Male | B1 | 5.2 | – | 2505 | – |
| 63 | Male | A2 | – | 17.8 | 2079 | 40.8 |
| 64 | Male | A2 | 3865 | 19.02 | Undetectable | 22.54 |
| 65 | Male | B2 | 5.77 | 16.3 | Undetectable | 34.1 |
| 66 | Male | C3 | 3.38 | 5.5 | 4.54 | 7.4 |
| 67 | Male | N3 | 3991 | 7.2 | 2041 | 9.1 |

| | | | | | | |
|---|---|---|---|---|---|---|
| 68 | Male | A3 | 5929 | 5.4 | Undetectable | 20.9 |
| 69 | Male | C3 | 5.38 | 13.6 | 3.25 | 15 |
| 70 | Male | N | 6491 | – | 6253 | 40.39 |
| 71 | Male | N2 | 4575 | 15.87 | – | – |

**Table S2**. PubMed-based compilation of IRIS-related infectious events in children and adolescents infected with HIV (January 1st, 1979 through August 30th, 2018).

| | | | | | | | Before ART | | After ART | | |
|---|---|---|---|---|---|---|---|---|---|---|---|
| **Reference** | **Case** [a] | **Age** [b] | **Sex** | **IRIS-related infectious event** | **Country** | **ART timing** [c] | **Viral load** | **CD4 count** | **Viral load** | **CD4 count** | **Fatal outcome** |
| (Bonkowsky et al., 2002) | 1 | 12 | Male | Chronic vascular infection | USA | 20 | NI [d] | 50 | <50 | 257 | No |
| (Gamero et al., 2012) | 1 | 4 months | Male | BCGitis | Spanish | 3 | 10,000 | 696 | NI | NI | No |
| (Innes et al., 2009) | 2 | 8 months | Female | Tuberculosis | South Africa | 20 | 43,200 | 908 | Undetectable | 1,214 | No |
| (Koppel et al., 2010) | 1 | 4 months | Male | BCGitis | Germany | 8 | NI | 583 | NI | NI | NI |
| (Koppel et al., 2010) | 2 | 9 months | Female | BCGitis | Germany | 12 | NI | NI | NI | NI | NI |
| (Koppel et al., 2010) | 3 | NI | NI | BCGitis | Germany | 11 | NI | NI | NI | NI | NI |
| (Kroidl et al., 2006) | 1 | 18 months | Male | BCGitis | Germany | 20 | 354,607 | 23,000 | 708 | 322,000 | No |
| (Narendran et al., 2006) | 1 | 12 | Male | Tuberculosis | India | <1 | NI | 51,000 | NI | 225,000 | No |
| (Nuttall et al., 2004) | 1 | 12 | Male | Progressive multifocal leukoencephalopathy | South | 5 | 96,000 | 12,000 | Undetectable | 49,000 | No |

| | | | | | | | | | | | |
|---|---|---|---|---|---|---|---|---|---|---|---|
| | | | | | Africa | | | | | | |
| (Oberdorfer et al., 2009) | 1 | 9 | Male | Progressive multifocal leukoencephalopathy | Thailand | 14 | 185,976 | 4,000 | 3,220 | 30,000 | No |
| (Perez et al., 2009) | 1 | 11 | Male | Graves' disease | African-American | 120 | 550,175 | 1,000 | Undetectable | 689,000 | No |
| (Puthanakit et al., 2006a) | 1 | 6 | Female | Mycobacterium avium | Thailand | 2 | Log 5.56 | 4% | Log 1.70 | 5% | No |
| (Puthanakit et al., 2006a) | 2 | 7 | Male | Mycobacterium avium | Thailand | 2 | Log 5.47 | 1% | Log 2.60 | 4% | No |
| (Puthanakit et al., 2006a) | 3 | 7 | Male | Mycobacterium avium | Thailand | 5 | NI | 1% | Log 4.78 | 9% | No |
| (Puthanakit et al., 2006a) | 4 | 9 | Female | Mycobacterium avium and Herpes labialis | Thailand | 26 | Log 5.57 | 6% | Log 1.70 | 19% | Yes |
| (Puthanakit et al., 2006a) | 5 | 10 | Male | *Mycobacterium scrofulaceum* | Thailand | 2 | Log 5.02 | 0% | Log 1.70 | 2% | No |
| (Puthanakit et al., 2006a) | 6 | 11 | Male | *Mycobacterium scrofulaceum* | Thailand | 2 | Log 4.86 | 0% | NI | 7% | Yes |
| (Puthanakit et al., 2006a) | 7 | 9 | Female | *Mycobacterium scrofulaceum* | Thailand | 3 | Log 5.40 | 2% | Log 2.22 | 8% | No |
| (Puthanakit et al., 2006a) | 8 | 8 | Male | *Mycobacterium kansasii* | Thailand | 3 | Log 5.50 | 0% | Log 2.60 | 5% | No |
| (Puthanakit et al., 2006a) | 9 | 7 | Male | *Mycobacterium simiae* | Thailand | 18 | NI | 0% | NI | 17% | No |
| (Puthanakit et al., 2006a) | 10 | 5 | Female | *Mycobacterium tuberculosis* | Thailand | 12 | Log 5.88 | 2% | Log 1.70 | 6% | No |
| (Puthanakit et al., 2006a) | 11 | 12 | Male | *Mycobacterium tuberculosis* | Thailand | 16 | NI | 4% | Log 1.70 | 14% | No |
| (Puthanakit et al., 2006a) | 12 | 3 | Male | *Mycobacterium tuberculosis* | Thailand | 31 | Log 5.88 | 3% | Log 2.09 | 11% | No |

| | | | | | | | | | | | |
|---|---|---|---|---|---|---|---|---|---|---|---|
| (Puthanakit et al., 2006a) | 13 | 8 | Female | BCGitis | Thail and | 3 | Log 4.38 | 3% | Log 1.87 | 6% | No |
| (Puthanakit et al., 2006a) | 14 | 8 | Female | BCGitis | Thail and | 4 | Log 5.88 | 0% | Log 2.21 | 2% | No |
| (Puthanakit et al., 2006a) | 15 | 6 | Male | Dermatomal Varicella-Zoster Virus | Thail and | 2 | Log 5.22 | 1% | Log 2.58 | 3% | No |
| (Puthanakit et al., 2006a) | 16 | 6 | Female | Dermatomal Varicella-Zoster Virus | Thail and | 2 | Log 5.33 | 3% | Log 2.45 | 10% | No |
| (Puthanakit et al., 2006a) | 17 | 14 | Female | Dermatomal Varicella-Zoster Virus | Thail and | 4 | Log 4.82 | 2% | Log 1.70 | 5% | No |
| (Puthanakit et al., 2006a) | 18 | 6 | Female | Dermatomal Varicella-Zoster Virus | Thail and | 6 | Log 5.65 | 3% | Log 2.36 | 8% | No |
| (Puthanakit et al., 2006a) | 19 | 10 | Female | Dermatomal Varicella-Zoster Virus | Thail and | 15 | NI | 0% | NI | 8% | No |
| (Puthanakit et al., 2006a) | 20 | 6 | Female | Dermatomal Varicella-Zoster Virus | Thail and | 21 | Log 4.82 | 2% | Log 2.39 | 6% | No |
| (Puthanakit et al., 2006a) | 21 | 5 | Female | HSV labialis | Thail and | 2 | Log 5.24 | 6% | Log 2.83 | 17% | No |
| (Puthanakit et al., 2006a) | 22 | 10 | Female | HSV labialis | Thail and | 4 | Log 5.43 | 15% | Log 1.88 | 14% | No |
| (Puthanakit et al., 2006a) | 23 | 11 | Male | HSV labialis | Thail and | 18 | Log 5.88 | 0% | Log 2.63 | 7% | No |
| (Puthanakit et al., 2006a) | 24 | 7 | Female | HSV labialis | Thail and | 31 | Log 5.88 | 2% | Log 2.20 | 15% | No |
| (Puthanakit et al., 2006a) | 25 | 11 | Female | HSV encephalitis | Thail and | 18 | Log 5.09 | 2% | Log 1.70 | 3% | Yes |
| (Puthanakit et al., 2006a) | 26 | 8 | Male | Cryptococcal meningitis | Thail and | 2 | Log 5.17 | 12% | Log 2.19 | 19% | No |
| (Puthanakit et al., 2006a) | 27 | 15 | Female | Cryptococcal meningitis | Thail and | 2 | Log 5.76 | 5% | Log 2.56 | 11% | No |
| (Puthanakit et al., 2006a) | 28 | 10 | Female | Cryptococcal meningitis | Thail and | 7 | Log 4.80 | 1% | Log 1.70 | 10% | No |

| | | | | | | | | | | | |
|---|---|---|---|---|---|---|---|---|---|---|---|
| (Puthanakit et al., 2006a) | 29 | 2 | Female | Guillain-Barré syndrome | Thailand | 3 | Log 5.88 | 12% | Log 3.48 | 26% | No |
| (Puthanakit et al., 2007) | 7 | 3 | Male | Cryptococcal meningitis | Thailand | 7 | Log 5.33 | 1% | NI | NI | Yes |
| (Puthanakit et al., 2007) | 8 | 6 months | Male | CMV retinitis and colitis | Thailand | 9 | Log 5.88 | 11% | NI | NI | Yes |
| (Puthanakit et al., 2007) | 11 | 11 | Female | CMV retinitis | Thailand | 22 | Log 5.09 | 2% | NI | NI | Yes |
| (Puthanakit et al., 2007) | 12 | 9 | Female | *Mycobacterium avium* complex | Thailand | 9.5 | Log 5.02 | 0% | NI | NI | Yes |
| (Wang et al., 2009) | 1 | 5 | Male | Varicella Zoster Virus | Peru | 2 | Log 5.90 | 346 (10%) | Log <2.60 | 770 (14%) | No |
| (Wang et al., 2009) | 2 | 7 | Female | Varicella Zoster Virus | Peru | 2 | Log 5.28 | 318 (14%) | Log <2.60 | 457 (22%) | No |
| (Wang et al., 2009) | 3 | 10 | Male | Varicella Zoster Virus | Peru | 6 | Log 6.45 | 2 (0%) | Log 4.81 | 140 (5%) | No |
| (Wang et al., 2009) | 4 | 9 | Female | Varicella Zoster Virus | Peru | 7 | Log 5.29 | 9 (1%) | Log 4.94 | 165 (8%) | No |
| (Wang et al., 2009) | 5 | 5 | Female | Varicella Zoster Virus | Peru | 11 | Log 4.52 | 334 (8%) | Log <2.60 | 539 (42%) | No |
| (Wang et al., 2009) | 6 | 9 | Female | Varicella Zoster Virus | Peru | 22 | Log 5.57 | 40 (6%) | Log <2.60 | 276 (15%) | No |
| (Wang et al., 2009) | 7 | 8 months | Female | *Mycobacterium tuberculosis* | Peru | 2 | Log 5.88 | 158 (7%) | Log 5.55 | 460 (7%) | No |
| (Wang et al., 2009) | 8 | 10 months | Male | *Mycobacterium tuberculosis* | Peru | 5 | Log 6.05 | 297 (2%) | Log 3.00 | 2,000 (15%) | No |
| (Wang et al., 2009) | 9 | 10 | Male | *Mycobacterium tuberculosis* | Peru | 6 | Log 5.20 | 59 (2%) | Log <2.60 | 391 (15%) | No |
| (Wang et al., 2009) | 10 | 1 | Male | *Mycobacterium tuberculosis* | Peru | 6 | Log 5.75 | 826 (18%) | Log 3.91 | 1,063 (11%) | No |

| | | | | | | | | | | | |
|---|---|---|---|---|---|---|---|---|---|---|---|
| (Wang et al., 2009) | 11 | 11 | Female | *Mycobacterium tuberculosis* | Peru | 28 | Log 6.26 | 40 (1%) | Log 5.45 | 1,226 (17%) | No |
| (Wang et al., 2009) | 12 | 1 | Female | BCGitis | Peru | 32 | Log 5.97 | 69 (2%) | Log 4.68 | 900 (14%) | No |
| (Wang et al., 2009) | 13 | 10 | Male | HSV labialis | Peru | 2 | Log 3.76 | 2 (0%) | Log <2.60 | 10 (1%) | No |
| (Wang et al., 2009) | 14 | 6 | Female | HSV labialis | Peru | 6 | Log 5.70 | 43 (2%) | Log 4.19 | 128 (3%) | No |
| (Wang et al., 2009) | 15 | 6 | Female | HSV labialis | Peru | 27 | Log 5.72 | 55 (1%) | Log <2.60 | 291 (11%) | No |
| (Wang et al., 2009) | 16 | 6 | Female | HSV labialis | Peru | 28 | Log 5.35 | 103 (8%) | Log 4.74 | 285 (12%) | No |
| (Wang et al., 2009) | 17 | 3 | Male | HSV labialis | Peru | 30 | Log 4.95 | 1591 (14%) | Log 3.80 | 1,733 (24%) | No |
| (Wang et al., 2009) | 18 | 4 | Male | HSV labialis | Peru | 34 | Log 5.74 | 4 (0%) | Log 4.05 | 248 (5%) | No |
| (Puthanakit et al., 2006b) | 1 | 10 | Male | Nontuberculous mycobacterial disease | Thail and | 2 | Log 5.02 | 0% | Log 1.7 | 2% | Yes |
| (Puthanakit et al., 2006b) | 2 | 11 | Male | Nontuberculous mycobacterial disease | Thail and | 2 | Log 4.86 | 0% | NI | 7% | Yes |
| (Puthanakit et al., 2006b) | 3 | 6 | Female | Nontuberculous mycobacterial disease | Thail and | 2 | Log 5.56 | 4% | Log 1.70 | 5% | No |
| (Puthanakit et al., 2006b) | 4 | 8 | Male | Nontuberculous mycobacterial disease | Thail and | 3 | Log 5.50 | 0% | Log 2.60 | 5% | No |
| (Puthanakit et al., 2006b) | 5 | 9 | Female | Nontuberculous mycobacterial disease | Thail and | 3 | Log 5.40 | 2% | Log 2.22 | 8% | No |
| (Puthanakit et al., 2006b) | 6 | 7 | Male | Nontuberculous mycobacterial disease | Thail and | 5 | NI | 1% | Log 4.78 | 9% | No |
| (Puthanakit et al., 2006b) | 7 | 9 | Female | Nontuberculous mycobacterial disease | Thail and | 26 | Log 5.57 | 6% | Log 1.70 | 19% | Yes |
| (Puthanakit et al., 2006b) | 8 | 7 | Male | Nontuberculous mycobacterial disease | Thail and | 10 | Log 5.47 | 1% | Log 2.60 | 4% | No |
| (Puthanakit et al., 2006b) | 9 | 7 | Male | Nontuberculous mycobacterial disease | Thail and | 23 | NI | 0% | NI | 17% | No |

| | | | | | | | | | | | |
|---|---|---|---|---|---|---|---|---|---|---|---|
| (Siberry and Tessema, 2006) | 1 | 9 | Male | BCGitis | Ethiopia | 1 | 122,000 | 9.6% (582) | NI | 716 | No |
| (Hesseling et al., 2003) | 1 | 9 months | Female | BCGitis | South Africa | 3 | NI | NI | 43,000 | 10% | No |
| (Sharp and Mallon, 1998) | 1 | 8 months | Male | BCGitis | Australia | 2 | $4.4x10^6$ | 10% | $7.7x10^4$ | 26% | No |
| (Viani, 2002) | 1 | 13 | Male | Sarcoidosis | USA | 36 | 750,000 | 5 cells/uL | <25 copies/mL | 441 cells/mL | No |
| (Sudjaritruk et al., 2012) | 1 | 14 | Female | *Penicillium marneffei* | Thailand | 8 | NI | 7.2% (39 cell/mm3) | <50 copies/mL | 11% (51 cells/mL) | No |
| (van Toorn et al., 2005) | 1 | 21 months | Female | Opsoclonus-myoclonus | South Africa | 2 | NI | 249 (3.5%) | 7,702 | 1,055 | Yes |
| (van Toorn et al., 2012) | 1 | 10 | Female | Neurotuberculosis | South Africa | 1 | NI | 55 (7%) | NI | 171 (11.7%) | Yes |
| (van Toorn et al., 2012) | 2 | 12 | Male | Neurotuberculosis | South Africa | 1 | NI | 274 | NI | 90 (8.3%) | No |
| (van Toorn et al., 2012) | 3 | 14 | Female | Neurotuberculosis | South Africa | 1 | NI | NI | 27 copies/mL | 61 (11.2%) | No |

| | | | | | | | | | | | |
|---|---|---|---|---|---|---|---|---|---|---|---|
| (van Toorn et al., 2012) | 4 | 21 months | Female | Neurotuberculosis | South Africa | 2 | NI | NI | NI | 1,496 (33%) | No |
| (Steenhoff et al., 2007) | 1 | 13 | Male | Cutaneous *Mycobacterium avium* | USA | 8 | >100,000 copies/mL | 1% | <50 copies/mL | 8% | No |
| (Sharland et al., 1998) | 1 | 7 | Female | *Pneumocystis carinii* pneumonia | England | NI | Log 5.4 | 233 cells/mL | NI | 388 cells/mL | No |
| (Sharland et al., 1998) | 2 | 7 | NI | Toxoplasmosis | England | NI | Log 5.9 | 92 cells/mL | NI | 349 cells/mL | No |
| (Rabie et al., 2010) | 1 | 3 | Male | *Mycobacterium tuberculosis* | England | 7 | Log 5.6 | 2.60% | NI | NI | No |
| (Puthanakit et al., 2005) | 1 | 9 | Female | BCGitis | Thailand | 4 | Log 5.88 | 0% | Log 2.21 | 2% | No |
| (Puthanakit et al., 2005) | 2 | 8 | Female | BCGitis | Thailand | 10 | Log 5.88 | 1% | Log 2.53 | 7% | No |
| (Puthanakit et al., 2005) | 3 | 8 | Female | BCGitis | Thailand | 4 | Log 4.38 | 3% | Log 1.87 | 6% | No |
| (Puthanakit et al., 2005) | 4 | 10 months | Female | BCGitis | Thailand | 8 | Log 5.88 | 13% | Log 3.73 | 24% | No |
| (Van der Linden et al., 2010) | 1 | 8 months | Female | *Hepatitis B* | South Africa | 6 | 690,000 copies/mL | 0.78% | 13,000 | 6.60% | Yes |
| (Kalk et al., 2013) | 1 | 6 | Male | *Mycobacterium tuberculosis* meningitis | South Africa | 4 | 20,000 copies/mL | 113 cells/mL (16.5%) | <25 copies/mL | 212 (10.1%) | Yes |

| | | | | | | | | | | | |
|---|---|---|---|---|---|---|---|---|---|---|---|
| (Fernandes et al., 2009; Fernandes and Medina-Acosta, 2010) | 1 | 5 months | Female | BCGitis | Brazil | 4 | 750,000 copies/mL | NI | 30000 copies/mL | 53.50% | No |
| (Fernandes et al., 2009; Fernandes and Medina-Acosta, 2010) | 2 | 7 months | Female | BCGitis | Brazil | 6 | 5,130,000 copies/mL | 13.30% | 1,410 copies/mL | 29.70% | No |
| (Fernandes et al., 2009; Fernandes and Medina-Acosta, 2010) | 3 | 6 months | Female | BCGitis | Brazil | 3 | 1,200,000 copies/mL | 27.20% | 11,400 copies/mL | 30% | No |
| (Hatherill and Flisher, 2009) | 1 | 9 | Female | Delirium | South Africa | 3 | NI | 9 cells/µL (0.08%) | NI | NI | No |
| (Hatherill and Flisher, 2009) | 2 | 9 | Female | Delirium | South Africa | NI | NI | 77 cell/µL (1.4%) | NI | NI | No |
| (Hatherill and Flisher, 2009) | 3 | 11 | Female | Delirium/Tuberculosis | South Africa | 4 | NI | 25 cell/µL (0.86%) | NI | 175 cell/µL | Yes |
| (de Carvalho et al., 2010) | 1 | 11 | Female | *Molluscum Contagiosum* | Brazil | 60 | Log 6.8 | 112 | Log 2.8 | 1,069 | No |
| (Hassan et al., 2015) | 1 | 9 | Male | *Cryptococcosis* | South Africa | 5 | 25,000 copies/mL | 4 cells/mm3 | NI | NI | No |

| | | | | | | | | | | | |
|---|---|---|---|---|---|---|---|---|---|---|---|
| (Hassan et al., 2015) | 2 | 6 | Male | *Cryptococcosis* | South Africa | 28 | 121,317 copies/mL | 42 cells/mm3 | 400 copies/mL | 755 cells/mm3 | No |
| (Hassan et al., 2015) | 3 | 11 | Female | *Cryptococcosis* | South Africa | 4 | NI | 135 cells/mm3 | 400 copies/mL | 208 cells/mm3 | No |
| (Hassan et al., 2015) | 4 | 7 | Male | *Cryptococcosis* | South Africa | 49 | NI | 10 cells/mm3 | 400 copies/mL | 145 cells/mm3 | No |
| (Iro et al., 2013) | 1 | 11 | Female | Varicella Zoster Virus | Botswana / UK | 4 | 250,000 copies/mL | 6,000 cell/mL | 404 copies/mL | 30,000/mL | No |
| (Lankisch et al., 2012) | 1 | 5 months | NI | Nephrotic syndrome | Germany | 4 | >20,000,000 copies/mL | 534 cells/µL (18%) | 61,872 copies/mL | 781 cells/µL | No |
| (Miranda-Choque et al., 2012) | 1 | 9 months | NI | BCGitis | Peru | 4 | NI | NI | NI | NI | No |
| (Miranda-Choque et al., 2012) | 2 | 13 months | NI | BCGitis | Peru | 4 | NI | NI | NI | NI | No |
| (Miranda-Choque et al., 2012) | 3 | 6 months | NI | BCGitis | Peru | 9 | NI | NI | NI | NI | No |
| (Miranda-Choque et al., 2012) | 4 | 4 months | NI | BCGitis | Peru | 3 | NI | NI | NI | NI | No |

| | | | | | | | | | | | |
|---|---|---|---|---|---|---|---|---|---|---|---|
| (Miranda-Choque et al., 2012) | 5 | 5 months | NI | BCGitis | Peru | 7 | NI | NI | NI | NI | No |
| (Miranda-Choque et al., 2012) | 6 | 7 months | NI | BCGitis | Peru | 4 | NI | NI | NI | NI | No |
| (Miranda-Choque et al., 2012) | 7 | 6 months | NI | BCGitis | Peru | 6 | NI | NI | NI | NI | No |
| (Miranda-Choque et al., 2012) | 8 | 16 months | NI | BCGitis | Peru | 4 | NI | NI | NI | NI | No |
| (Nandy and Shah, 2015) | 1 | 7 | Female | Giardiasis | NI | 4 | NI | 158 cells/mm3 | NI | NI | No |
| (Ramdial et al., 2011) | 1 | 9 | Male | *Cryptococcosis in kidney* | South Africa | 10 | 4,300 copies/mL | 225 cells/mm3 | <25 copies/mL | 419 cells/mm3 | No |
| (Ramdial et al., 2011) | 2 | 7 | Female | *Cryptococcosis in kidney* | South Africa | 15 | 3,300 copies/mL | 80 cells/mm3 | <25 copies/mL | 62 cells/mm3 | No |
| (Schwenk et al., 2014) | 1 | 15 | Female | Progressive multifocal leukoencephalopathy | NI | NI | 82,000copies/mL | 5 cells/mm3 | NI | NI | Yes |
| (Van Rie et al., 2016) | 1 | 10 months | NI | Abdominal + pulmonary tuberculosis | South Africa | 2 | 1,700,000 | 11% | NI | NI | No |
| (Van Rie et al., 2016) | 2 | 8 months | NI | Pulmonary tuberculosis | South Africa | 2 | 9,900,000 | 16% | NI | NI | No |

| | | | | | | | | | | | |
|---|---|---|---|---|---|---|---|---|---|---|---|
| (Van Rie et al., 2016) | 3 | 24 months | NI | Abdominal + pulmonary tuberculosis | South Africa | 9 | 63,000 | 7.90% | 400 | 15.60% | No |
| (Van Rie et al., 2016) | 4 | 81 months | NI | Abdominal + lymph node + pulmonary tuberculosis | South Africa | 2 | 481,202 | 29.10% | 5,608 | 33.60% | No |
| (Van Rie et al., 2016) | 5 | 8 months | NI | Pulmonary tuberculosis | South Africa | 4 | 690,000 | 50.50% | NI | NI | No |
| (Van Rie et al., 2016) | 6 | 53 months | NI | Pulmonary tuberculosis | South Africa | 5 | 230,000 | 20.60% | NI | NI | No |
| (Van Rie et al., 2016) | 7 | 30 months | NI | Pulmonary tuberculosis | South Africa | 2 | 380,000 | 24.10% | NI | NI | No |
| (Pereira et al., 2016) | 1 | 13 | Female | Opsoclonus-Myoclonus-Ataxia | India | 60 | 28,387 | 77 cell/mL | 165 | 429 cell/mL | No |
| (Dunkley-Thompson et al., 2008) | 1 | 7 months | Male | BCGitis | Jamaica | 21 | NI | 40% | NI | 40.70% | No |
| (Dunkley-Thompson et al., 2008) | 2 | 7 months | Male | BCGitis | Jamaica | 42 | NI | 16% | NI | 43.70% | No |
| (Dunkley-Thompson et al., 2008) | 3 | 9 months | Female | BCGitis | Jamaica | 28 | NI | 43% | NI | 53.60% | No |

[a] Case number by reference
[b] Age in years; otherwise in months
[c] Days to onset of IRIS event after ART
[d] Not informed

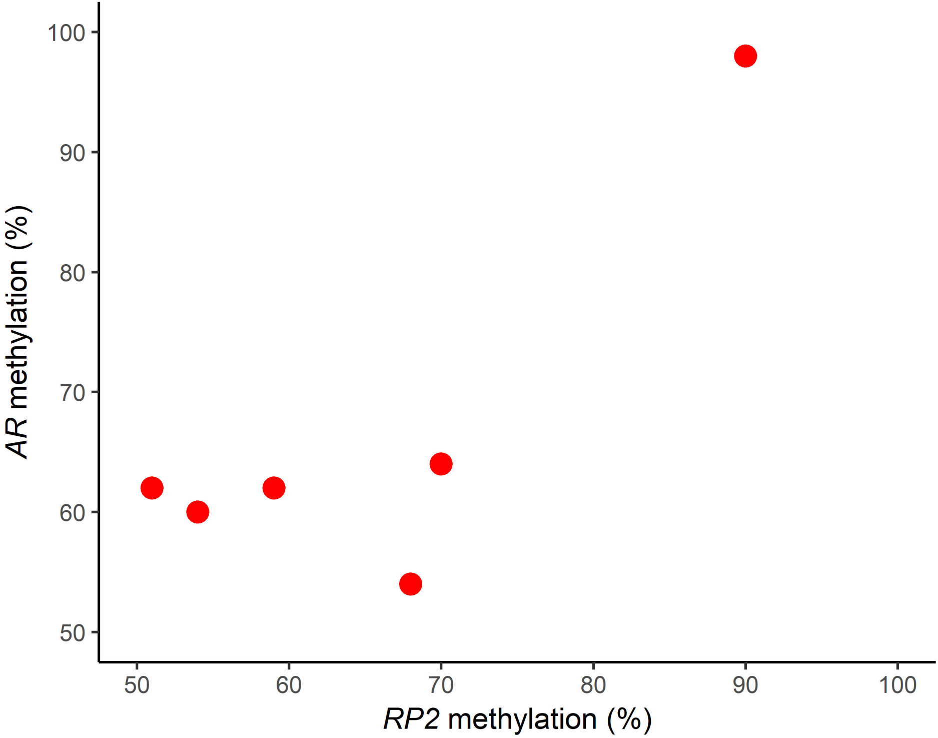
100
90
80
70
60
50
AR methylation (%)
50
60
70
80
90
100
RP2 methylation (%)
